# Safeguarding spermatogenesis from retrotransposon insertions by forming ecDNA

**DOI:** 10.1101/2025.05.11.653319

**Authors:** Lauren Tracy, ZZ Zhao Zhang

## Abstract

Retrotransposon mobilization in germline cells enables the rewriting of genetic information to drive genome innovation, species evolution, and adaptation through the generation of de novo mutations. However, uncontrolled mobilization can cause DNA breaks and genome instability, often leading to sterility. How germ cells balance retrotransposon-induced genome innovation with the need for genomic integrity remains poorly understood. Here, we used *Drosophila* spermatogenesis as a model to investigate retrotransposon mobilization dynamics. Although many retrotransposon families are transcriptionally active, we found that the LTR-retrotransposon *nomad* completes the full mobilization cascade—including mRNA export, protein translation, and reverse transcription—to produce double-stranded DNA (dsDNA) the most efficiently. Strikingly, despite successfully generating dsDNA, *nomad* rarely achieves genomic reintegration. Instead, its newly synthesized DNA predominantly forms extrachromosomal circular DNA (ecDNA). These findings suggest that ecDNA formation acts as a protective mechanism to sequester retrotransposon-derived DNA and prevent widespread genomic integration during spermatogenesis, thereby preserving genome stability while allowing limited retrotransposon activity.

## Introduction

Organisms evolve through changes in germline DNA, which can range from single-nucleotide variations to larger insertions and deletions spanning kilobases. One major source of large genomic insertions is transposable elements—segments of DNA capable of changing their position within the genome. These elements mobilize either by excising themselves and reinserting elsewhere or by copying themselves through an RNA intermediate before integration (Rubin et al. 1982; Boeke et al. 1985; Kazazian and Moran 2017; Fueyo et al. 2022). The latter category, retrotransposons, can significantly expand genome size since they generate new insertions while retaining their original copies. During this process, they reverse transcribe their mRNA into double-stranded DNA for making new genomic copies. As a result, transposable elements constitute approximately 45% of the human genome and 20% of the *Drosophila* genome (Lander et al. 2001; Merel et al. 2020).

Retrotransposon mobilization events in the germline can have profound and lasting impacts on host biology, as they introduce heritable genetic variation that may be either deleterious or beneficial. While some insertions are harmful—such as a LINE-1 insertion linked to hemophilia A in two patients (Kazazian et al. 1988)—others have been retained through evolution due to their advantageous effects. In *Drosophila*, transposable elements account for a substantial fraction of spontaneous mutations, many with negative consequences (Pasyukova and Nuzhdin 1993; Adrion et al. 2017). Yet across species, specific retrotransposon insertions have been co-opted for critical biological functions. In primates, for instance, retrotransposons contribute to placental development; notably, an envelope protein derived from an LTR-retrotransposon has been repurposed for the formation of syncytiotrophoblasts (Mi et al. 2000). LTR elements also provide regulatory sequences that influence gene expression during early mammal development (Macfarlan et al. 2012; Wang et al. 2014; Grow et al. 2015; Xiang et al. 2022). Similarly, in *Drosophila*, insertion of the *roo* retrotransposon upstream of a stress-response gene has been shown to enhance cold-stress tolerance (Merenciano et al. 2016). These examples highlight how retrotransposon activity in the germline can drive both evolutionary innovation and adaptive function, while also posing risks to genome stability.

Given the profound impact of germline transposon insertions, understanding how their mobilization is regulated is critical to balancing evolutionary innovation with genome integrity. Using *Drosophila* spermatogenesis as a model, we demonstrate that retrotransposons can be transcriptionally derepressed, leading to the production of mRNA capable of reverse transcription into double-stranded DNA. However, rather than integrating into the genome, these replicated DNA molecules are sequestered as extrachromosomal circular DNA (ecDNA), resulting in minimal, if any, insertion events. Our findings suggest that hosts safeguard their germline by redirecting retrotransposon-derived DNA into ecDNA, thereby preventing unchecked mobilization and potential genomic instability.

## Results

### Transposon activation during *Drosophila* spermatogenesis does not cause sterility

In germ cells, a small RNA pathway called the piRNA pathway controls transposon expression by slicing transposon transcripts in the cytoplasm and inducing heterochromatin formation around transposons to mitigate their activity at the transcriptional level (Cox et al. 1998; Cox et al. 2000; Aravin et al. 2006; Girard et al. 2006; Lau et al. 2006; Saito et al. 2006; Vagin et al. 2006; Brennecke et al. 2007). This pathway is in place to mitigate the effects of transposon activity on the germline genome, which is passed to the next generation. To further boost transposon activity during spermatogenesis, we silenced the repressive piRNA pathway by RNAi. Specifically, we used the maternal triple driver (MTD-Gal4) driver line to deplete the piRNA pathway components Aubergine (Aub) and Argonaute3 (Ago3) by RNAi. In the testes from 3-day-old flies, Aub and Ago3 are efficiently depleted at the RNA level and undetectable at the protein level by immunostaining (Supplemental Fig. 1). Notably, even with the strong suppression of the piRNA pathway, *Drosophila* spermatogenesis remains normal, and flies remains fertile with slightly lower fecundity (Fig. 1A, B).

**Figure 1.**
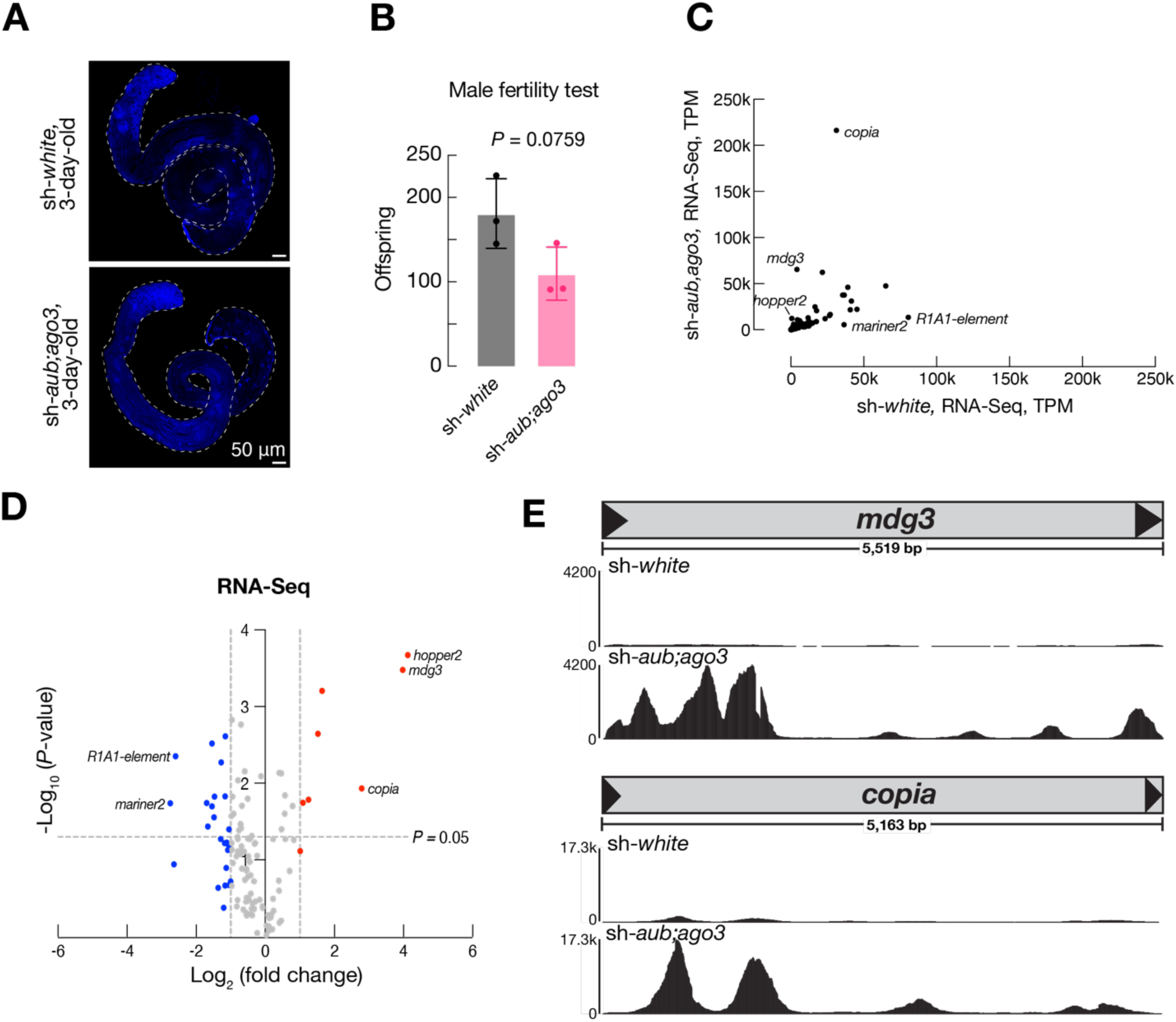
Disrupting the piRNA pathway in the male germline has minimal effects on fertility. **A.** Whole testes of 3-day-old sh-*white* and sh-*aub;ago3* flies. **B.** Offspring produced by sh-*white* or sh-*aub;ago3* males crossed with new female w^1118^ virgins over 6 days. 3 replicates per genotype. *P*-value determined by an unpaired t-test. **C.** Transposon expression in testes from 3-day-old sh-*white* and sh-*aub;ago3* flies. RNA sequencing data is from two replicates per genotype. **D.** Fold change in transposon RNA expression of sh-*white* vs. sh-*aub;ago3* testes from 3-day-old flies. Blue points, Log_2_(fold change) <-1. Red points, Log_2_(fold change) > 1. *P*-value determined by an unpaired t-test. **E.** RNA production across the length of *mdg3* or *copia* as visualized though IGV (Integrative Genomics Viewer).

To test whether transposons are indeed fully unleashed upon the piRNA pathway blockage, we surveyed the activation of transposons at the transcription level by performing RNA-Seq experiments. We observed drastic activation of some transposon families when the piRNA regulation is lifted. Among the 122 transposon families with detectable RNA expression in the testes, 8 increased their mRNA by more than 2-fold (Fig. 1D). Notably, among all families examined, including both DNA transposons and retrotransposons, LTR-retrotransposons are the ones that showed the most drastic changes: *copia* for 7-fold, *mdg3* for 16-fold, and *diver* for 3-fold (Fig. 1C-E). In summary, we found that silencing the piRNA pathway results in transposon activation during spermatogenesis but not animal sterility, indicating a protective program that safeguards germline genome from transposon mobilization.

### ecDNA production suggests accomplishment of reverse transcription

LTR-retrotransposons start their life cycle by having their RNA transcribed and transported to the cytoplasm to express the Gag and Pol proteins encoded by the transposon (Shiba and Saigo 1983; Garfinkel et al. 1985; Kazazian and Moran 2017; Wells and Feschotte 2020; Fueyo et al. 2022). Pol then works with host factors to reverse transcribe the LTR-retrotransposon RNA into a DNA copy for making new insertions (Boeke et al. 1985). Therefore, having mRNA transcribed does not guarantee the occurrence of downstream steps. Our lab previously found that LTR-retrotransposons use a DNA circularization step to complete second strand synthesis, and this step results in massive production of ecDNA (Yang et al. 2023). Therefore, to determine whether LTR-retrotransposons can accomplish downstream steps, we targeted the circular DNA form of these transposons through ecDNA sequencing on control (sh-*white*) and transposon derepressed (sh-*aub;ago3*) testes. Among the LTR-retrotransposons that significantly increased their mRNA, only two of them significantly increased ecDNA production: *copia* and *mdg3* (Fig. 2C, D). This indicates that mRNA expression is not a reliable indicator of downstream activities. Furthermore, we noticed the LTR-retrotransposon *nomad* dominantly manufactures the most ecDNA when transposons are derepressed. Specifically, *nomad* produces 448-fold more ecDNA in sh-*aub;ago3* testes compared to sh-*white* testes (Fig. 2A). From all transposon-derived ecDNA, *nomad* makes 68% of all circles. Meanwhile, we found that two LTR-retrotransposons (*297, 412*) could produce ecDNA at low levels even in control testes (Fig. 2B). And their ecDNA production significantly decreased when *nomad* and other transposons are unleashed for ecDNA production. This suggests that different retrotransposons compete with host factors for their DNA replication and ecDNA formation.

**Figure 2.**
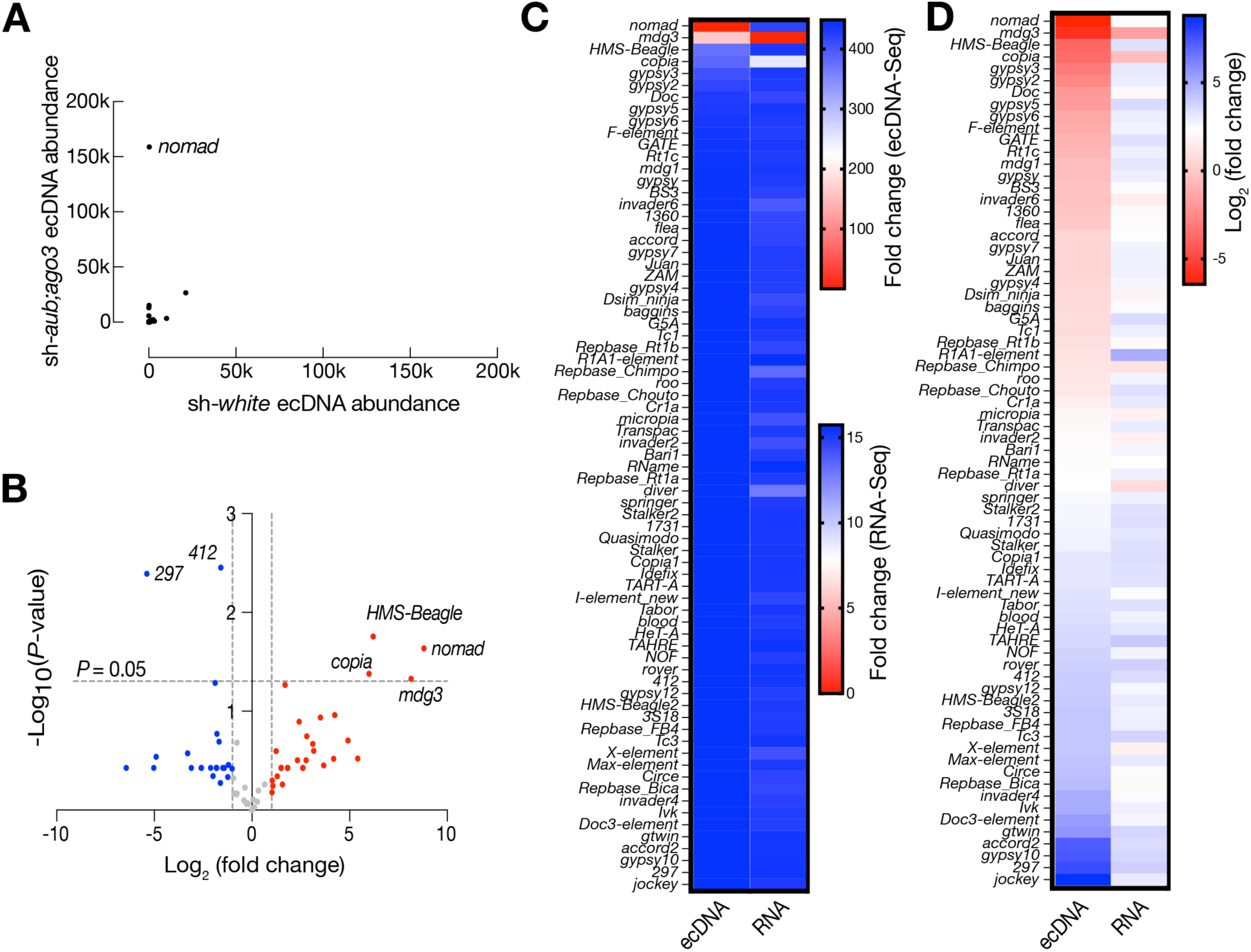
***nomad* abundantly produces ecDNA in the testis**. **A.** Transposon ecDNA abundance in the testes of sh-*white* and sh-*aub;ago3* 3-day-old flies. Data is from ecDNA sequencing of 3 biological replicates per genotype. Data is normalized by mitochondrial DNA abundance in each sample. **B.** Volcano plot of ecDNA detected from ecDNA sequencing in control vs. sh-*aub;ago3* tests. *P*-value determined by an unpaired t-test. Blue points, Log_2_(fold change) <-1. Red points, Log_2_(fold change) > 1. **C.** Heatmap comparing ecDNA abundance and RNA expression for all transposons with detectable ecDNA in sh-*white* vs sh-*aub;ago3* testes at 3 days old. **D.** Heatmap comparing ecDNA abundance and RNA expression for all transposons with detectable ecDNA in sh-*white* vs sh-*aub;ago3* testes at 3 days old, plotted using a Log_2_ scale.

To confirm the circular DNA sequencing results, we used divergent PCR to identify ecDNA production from testes. This method will only produce a PCR product if ecDNA is formed from retrotransposons. By divergent PCR, we validated that *nomad* ecDNA is abundantly produced in transposon-activated testes, and *HMS-Beagle* ecDNA are also produced, but at a lower abundance, consistent with our ecDNA sequencing data (Fig. 3A; Supplemental Fig. 3A). Additionally, we examined if *nomad* could achieve ecDNA production in a different genetic background with the piRNA pathway disrupted. As such, we examined *nomad* ecDNA production in *rhino* (*rhi*) mutants. Rhino is upstream of Aub and Ago3 in the piRNA pathway and helps initiate transcription of piRNAs for use in targeting and cleaving retrotransposon transcripts (Klattenhoff et al. 2009; Mohn et al. 2014; Zhang et al. 2014; Yu et al. 2018). In *rhino* mutant flies, *nomad* also abundantly produces ecDNA as examined by divergent PCR (Supplemental Fig. 3B). Taken together, our data reveal that *nomad* is the most active retrotransposon to accomplish the downstream activation process during *Drosophila* spermatogenesis.

**Figure 3.**
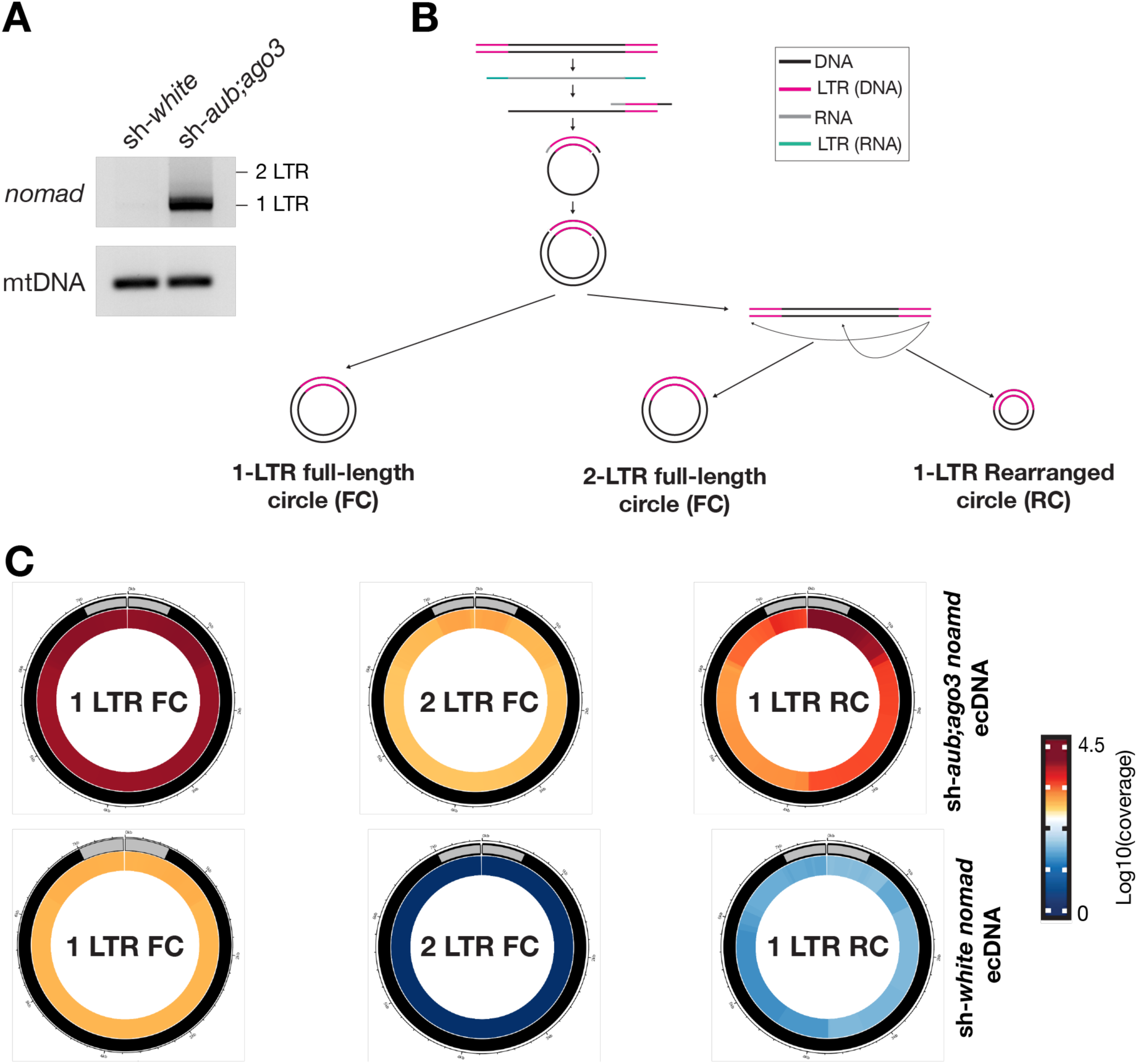
*nomad* predominantly makes 1-LTR ecDNA. **A.** Divergent PCR for *nomad* ecDNA. Input for PCR was DNA from 3-day-old testes with linear DNA digested. **B.** Model of how different forms or ecDNA are formed. 1-LTR circles are formed by ligation immediately following second-strand synthesis. 2-LTR circles are formed following second-strand synthesis and linearization of the cDNA. Then the linear DNA is then ligated together. 1-LTR partial circles likely be formed by self-integration. **C.** Circos plots of different types of *nomad* DNA circles and their abundance in testes from 3-day-old flies. Gray boxes signify the LTR region of *nomad*. FC = full-length circle. RC = rearranged circle. Circos plots are from representative replicated of the ecDNA sequencing experiment.

### LTR-retrotransposons manufacture distinct types of ecDNA

LTR-retrotransposons manufacture distinct types of ecDNA via different mechanisms (Fig. 3A). Often, the majority of the ecDNA produced during the 2^nd^-strand DNA synthesis is one LTR followed by the full internal sequence of the transposon (1-LTR full-length circles). Additionally, there are ecDNA circles that include the full transposon sequence (2-LTR full-length circles). This type of circle is likely formed by joining the two ends of the fully replicated double-stranded DNA precursors, indicating the completion of 2^nd^-strand DNA synthesis. Meanwhile, ecDNA can be produced from fragmented retrotransposon sequences by having one intact LTR (1-LTR rearranged). These circles can be produced when the integrase encoded from transposons uses the free end of the replicated double-stranded precursors to attack the internal sequences, a process known as “auto-integration”. Finally, fragmented retrotransposon sequences can form ecDNA by not having intact LTR (0-LTR rearranged) at low abundance, suggesting a direct ligation of degraded retrotransposon replicated DNA.

To explore the potential contribution of different programs on ecDNA production, we mined our ecDNA sequencing dataset to survey each circle type. When unleashed during spermatogenesis, 1-LTR full-length *nomad* circles represented 43.9% of the circles (Fig. 3B). We also detected 2-LTR full-length circles which represented 2.0% of *nomad* ecDNA, suggesting the completion of the 2^nd^-stranded DNA synthesis for some *nomad* precursors. Meanwhile, we detected 52.5% of *nomad* ecDNA as 1-LTR rearranged. This indicates that *nomad* can translate active integrase to potentially mediate the integration step. Finally, we observed 2.3% of *nomad* ecDNA as 0-LTR rearranged.

By expanding our analysis to other retrotransposon families, we learned that all LTR-retrotransposons can produce these different circle types. Consistently, the most abundant circles for each transposon contain a single intact LTR (*copia*: 73.9% 1-LTR full-length circles, 11.3% 1-LTR rearranged circles; *HMS-Beagle*: 30.2% 1 LTR full-length circles, 63.0% 1-LTR rearranged circles; *mdg3*: 19.0% 1-LTR full-length circles, 76.4% 1-LTR rearranged circles) (Supplemental Fig. 4D). The abundance of 1-LTR rearranged circles suggests that they possess active integrase for making new insertions. For *copia*, the 2-LTR full-length circles are the 2^nd^ most abundant ecDNA type (12.1%), while 1-LTR rearranged circles only take 11.3%. This indicates that more of the replicated full-length double-stranded precursors are used for end-end joining to form 2-LTR full-length circles, and that *copia* may possess weak integration activity.

### *nomad* activation in early spermatocytes

Since *nomad* is the most active transposon at generating ecDNA, we sought to use *nomad* as an example to investigate the retrotransposon activation process during spermatogenesis. Spermatogenesis is the process when a germline stem cell divides and undergoes meiosis at the spermatogonia stage, then enlarges and turns on transcription highly as a spermatocyte, then undergoes two rounds of meiosis to become a spermatid and eventually undergoes conformational changes to become mature sperm (Fabian and Brill 2012). Using whole testes as material for RNA-Seq, we found that *nomad* RNA expression does not significantly change in sh-*aub;ago3* testes compared to sh-*white* testes, further suggesting that RNA level is not a reliable predictor of a transposon’s ability for replication (Fig. 2C, D). To get an idea of when *nomad* may be active and how this activation is achieved, we used RNA FISH and immunofluorescence staining to visualize where *nomad* RNA and proteins are made during spermatogenesis.

In both in sh-*aub;ago3* and sh-*white* testes, *nomad* is expressed in spermatocytes (Fig. 4A, B). Notably, in the early spermatocytes, there is a difference in localization of the *nomad* RNA. While *nomad* RNA is restricted to only the nuclei in sh-*white* spermatocytes, in the testes with the piRNA pathway depleted, *nomad* RNA is also able to localize to the cytoplasm (Fig. 4B). This suggests that when the piRNA pathway is suppressed, *nomad* RNA is able to be exported into the cytoplasm for protein translation and initiation of the downstream steps of life cycle. Interestingly, *copia* RNA was not detectable when piRNA pathway is still in action. However, it is expressed in spermatogonia and spermatocytes in the testes upon depletion of the piRNA pathway, but the RNA is restricted to the nuclei of those cells (Fig. 4C). This could explain why we observed an increase in *copia* RNA, but less dramatic increase in *copia* ecDNA if the RNA is not making it to the cytoplasm (Fig. 2C, D). Together, our data suggest that piRNA pathway can suppress transposon activity at both transcriptional (*copia*) and post-transcriptional (*nomad*) levels.

**Figure 4.**
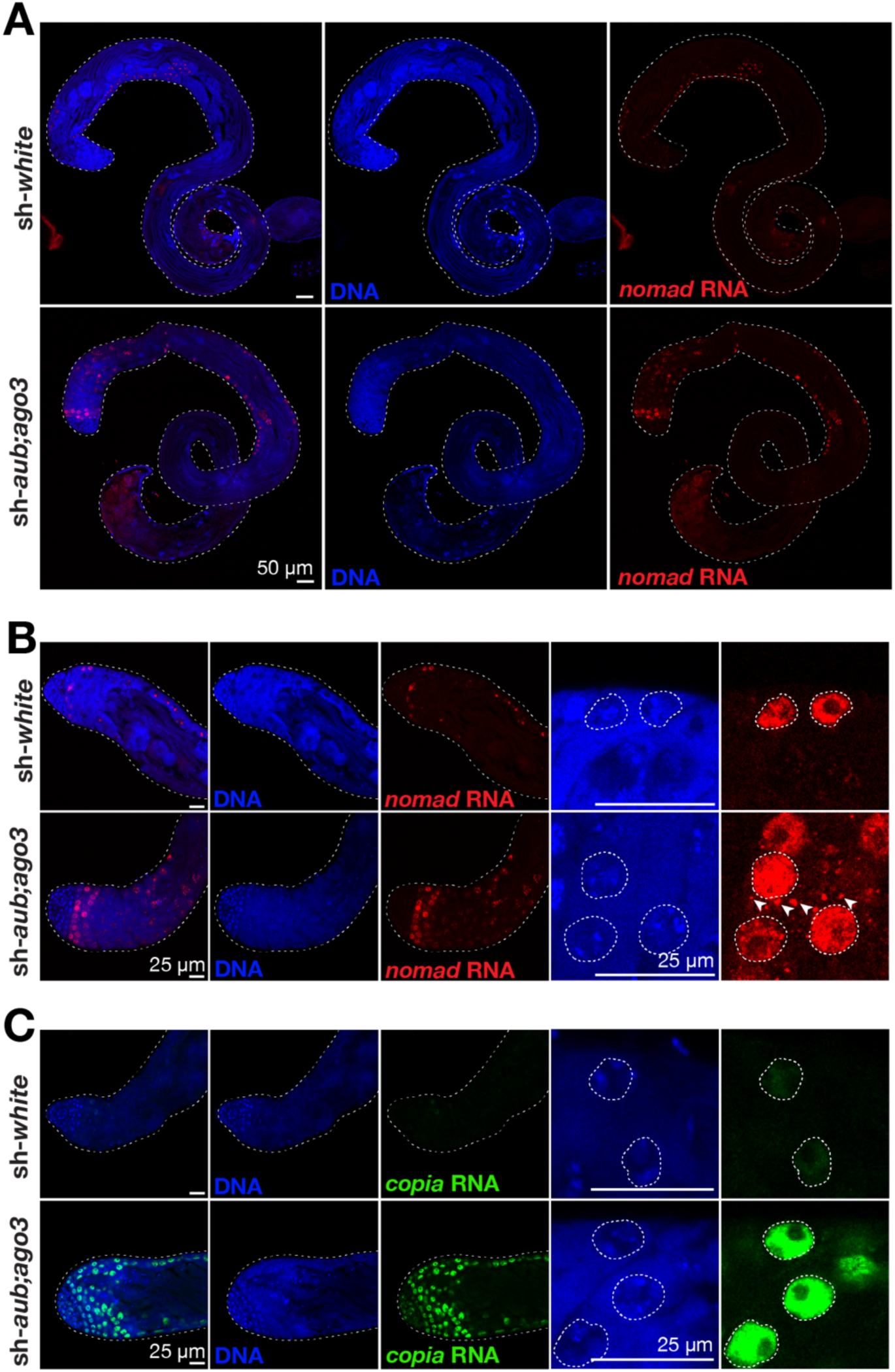
Efficient exportation of nomad, but not copia RNA, is correlated with increased ecDNA production. **A.** RNA FISH for *nomad* in the whole testis of 3-day-old flies. **B.** RNA FISH for detection of *nomad* RNA at the apical tip of the testis from 3-day-old flies. Arrows point to *nomad* RNA puncta in the cytoplasm. **C.** RNA FISH to detect *copia* RNA at the apical tip of the testis from 3-day-old flies.

As an LTR-retrotransposon, *nomad* encodes Gag and Pol proteins which are used for its lifecycle and also encodes a truncated *Env* protein (Whalen and Grigliatti 1998). To determine whether blocking piRNA pathway leads to *nomad* protein production, we raised a *nomad* Gag antibody. To examine the cell types that express Gag proteins, we next performed immunostaining experiments. While the sh-*white* testes only gave signals as background, in sh-*aub;ago3* testes from 3-day-old flies there is *nomad* Gag protein detected in early spermatocytes, the same cells that express *nomad* RNA (Supplemental Fig. 5A). As the flies age, there is an increase in *nomad* Gag accumulation in the spermatocytes in testes from 1-week-old flies compared to the 3-day-old testes (Supplemental Fig. 5B). Together, these experiments validated that *nomad* is suppressed at the post-transcriptional level by the piRNA pathway, and that *nomad* activation can lead to protein production to initiate downstream activation steps.

### Transposon integration is rare in the testes

The final step of the retrotransposon life cycle is integration of the new DNA copy of the transposon into a new location in the genome. While retrotransposons need to accomplish multiple stages of their life cycle—from transcription to reverse transcription—before being able to integrate, our ecDNA sequencing data indicate certain LTR-retrotransposon families are capable of achieving it. To look for new transposon integration events, we sequenced genomic DNA from sh-*white* and sh-*aub;ago3* testes of 1-week-old flies. As a control, we also sequenced the carcass of sh-*white* and sh-*aub;ago3* flies to provide a basis for which transposon insertions are already present in the genome of each genotype of flies (non-reference copies) (Fig. 5A). To avoid the potential false-positive events caused by the ligation step during library preparation, we used Nanopore tagmentation-based sequencing method, which bypasses the ligation step, to capture potential new insertion events.

**Figure 5.**
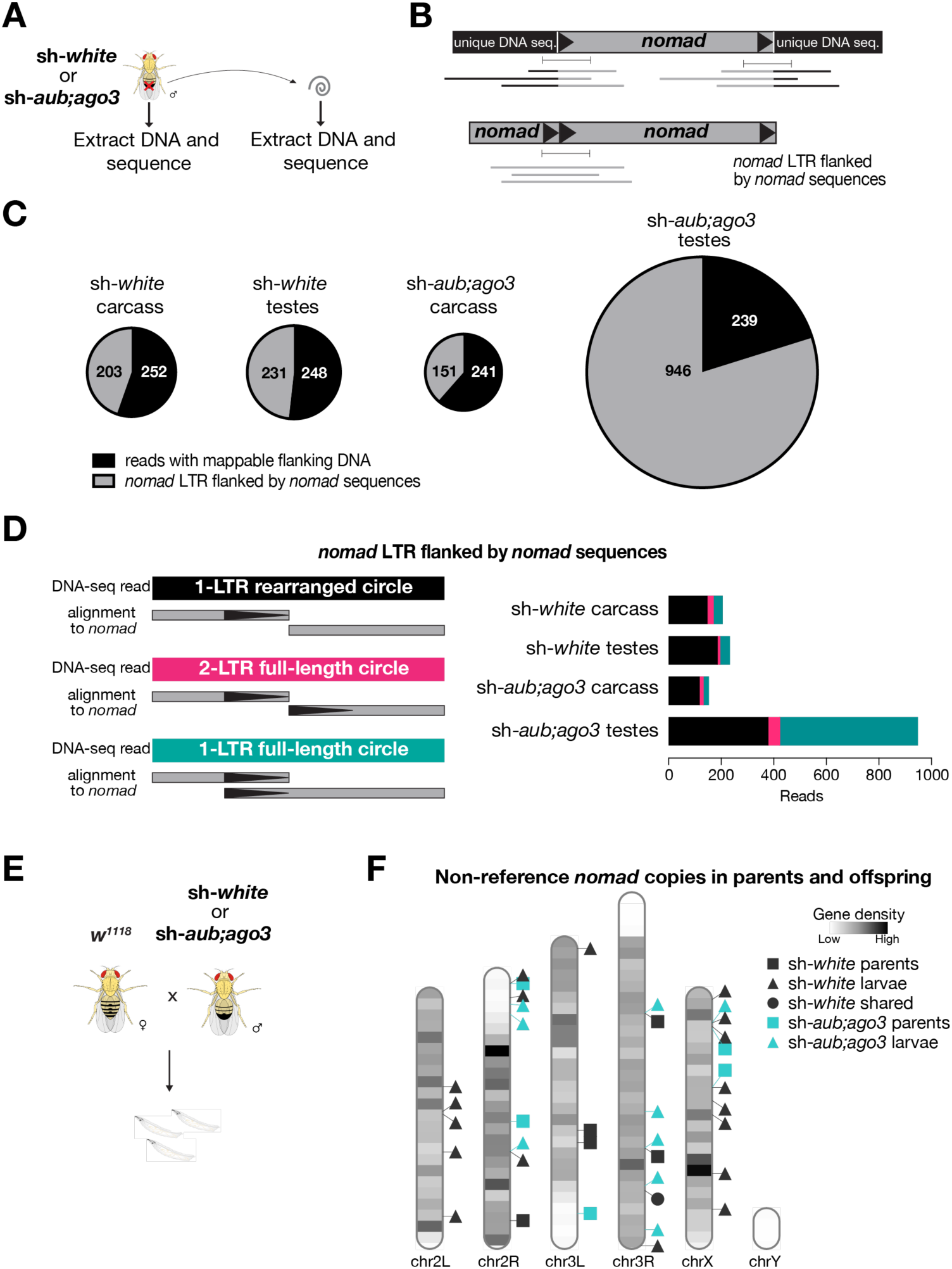
Transposition is rare in testes. **A.** Graphical representation of the genomic DNA sequencing experiment to look for transposon insertions in *Drosophila* testes compared to the carcass (without testes) of the fly. **B.** Representative model of one end *nomad* reads used in the analysis for panels C and D. **C.** Pie charts showing the type of DNA that is flanking the one end *nomad* reads from the carcass and testes of each genotype. One end reads that have flanking *nomad* sequences are either from *nomad* ecDNA or nested *nomad* insertions. **D.** Composition of different types of one-end *nomad* reads that only have *nomad* flanking sequence. The 1-LTR rearranged reads map to one end of *nomad* flanked by internal nomad sequence. The 2-LTR full-length circle reads map to both ends of *nomad* with 0 bp overlap between the two mapped regions. The 1-LTR full-length circle reads map to each end of *nomad* with overlap at the LTR region of *nomad*. **E.** Scheme for examining transposition in the offspring of males with transposons derepressed in the germline (panel F). **F.** Ideogram of non-reference *nomad* copies detected in the parents and offspring from crosses of wild-type (*w^1118^*) females crossed with either a control male (MTD-Gal4>sh-*white*) or male with transposons derepressed (MTD-Gal4>sh-*aub;ago3*) in the germline.

To first quantify the non-reference copies of *nomad*, we examined the insertion events that are supported by multiple sequencing reads and coordinated flanking sites from both ends. With 10-fold genome coverage, we obtained 0 copies of non-reference *nomad* copies from sh-*white* and only one from sh-*aub;ago3* testes. Notably, this *nomad* copy cannot be identified from carcass samples, suggesting polymorphisms of individual flies within the strains we used (Supplemental Fig. 6). Given that the landing sites for de novo insertions likely vary significantly among cells and we sequenced DNA from the whole testis, we reasoned that we would capture de novo insertions by only having one supporting read that flanks one end of the transposons (Fig. 5B). By performing such analysis, we identified ∼240 reads are flanked by mappable unique DNA sequences from all samples (Fig. 5C). Given that these read numbers were similar between sh-*white* and sh-*aub;ago3* testes, these events were unlikely caused by *nomad* hyper-activation when the piRNA pathway is blocked.

Strikingly, we identified 946 reads that are flanked by *nomad* sequences from sh *aub;ago3* testis (vs 150-203 reads in other control samples) (Fig. 5C), suggesting that the tagmentation method captures a fragment of *nomad* ecDNA for sequencing. The majority of these reads (55.3%) are able to map to each end of *nomad*, with an overlap at the LTR region (Fig. 5D). These reads are most likely from 1-LTR full-length ecDNA circles. Only 4.8% of the one-end *nomad* reads were immediately flanked by the following LTR with 0 bp overlap, indicating that these reads likely arise from 2-LTR circles (Fig. 5D). Lastly, 35.6% of the reads contained 1 LTR followed by internal *nomad* sequence. These reads could be from 1-LTR rearranged circles or potentially nested *nomad* insertions in the genome (Fig. 5D). Accordingly, these potential 1-LTR rearranged or nested insertion reads were the most abundant type in sh-*white* carcass (71.4%), sh-*white* testes (79.7%), and sh-*aub;ago3* carcass (76.8%) (Fig. 5D). Without ecDNA enrichment, capturing this large amount of ecDNA from whole testis DNA sequencing suggests that the majority of the replicated *nomad* molecules form ecDNA, but rarely achieve the integration step, which would cause DNA damage. Consistently, we did not observe difference of γ-H2Av signals––a hallmark for DNA breaks––in the sh-*aub;ago3* testes compared to the sh-*white* testes (Supplemental Fig. 7).

De novo insertions that occur during spermatogenesis can be transmitted to offspring and retained as non-referenced copies in all cells of the resulting animals. To further validate our finding that retrotransposons fail to generate new insertions during spermatogenesis, we crossed *sh-white* and *sh-aub;ago3* males with *w^1118^* females and sequenced DNA from their offspring at the larval stage (Fig. 5E). With 11-fold genome coverage, we identified 19 non-referenced copies of *nomad* in the progeny of *sh-white* males and 10 copies in the progeny of *sh-aub;ago3* males (Fig. 5F). Beyond *nomad*, we surveyed the new insertion events in both testes and progenies for other retrotransposons. We found similar as *nomad*, other retrotransposons, *copia*, *HMS-Beagle*, and *mdg3* all failed to achieve new integration events when the piRNA pathway is blocked (Supplemental Fig. 6; Supplemental Fig. 8). The absence of new insertion events in both testes and offspring, despite the presence of hyper transposon activity in the testes of these males, suggests that the formation of extrachromosomal circular DNA (ecDNA) serves to protect the germline from transposon invasion, thereby safeguarding genomic integrity during spermatogenesis.

## Discussion

In this study, we reveal that *Drosophila* spermatogenesis employs a unique strategy to protect the germline genome from retrotransposon insertions. While multiple retrotransposon families are transcriptionally activated when the piRNA pathway is depleted, we find that only a few, most notably the LTR-retrotransposon *nomad*, progress through the entire retrotransposon life cycle. Strikingly, instead of integrating into the genome, *nomad* DNA intermediates are predominantly converted into ecDNA. Our genomic analysis of testes and progeny validated that new insertions are rare, even when retrotransposon activity is elevated, suggesting that ecDNA formation acts as a critical layer of genome protection during spermatogenesis.

Our findings highlight a multilayered regulatory framework that controls retrotransposon mobilization. At the transcriptional level, the piRNA pathway silences many transposons by promoting heterochromatin formation (Brower-Toland et al. 2007; Rangan et al. 2011; Sienski et al. 2012). However, transcription alone is not predictive of downstream activity. For instance, while *copia* massively transcribes mRNAs upon the loss of piRNA protection, they remain confined to the nucleus and only minimally proceed to DNA synthesis. In contrast, *nomad* RNA is exported to the cytoplasm, translated into Gag proteins, and reverse transcribed into double-stranded DNA. This suggests that post-transcriptional regulation—such as nuclear retention of RNA or inhibition of protein production—represents an important control point in addition to transcriptional silencing.

Beyond translation, additional post-replication mechanisms shape retrotransposon outcomes. Our data show that even when reverse transcription occurs, most resulting DNA does not integrate into the genome. Instead, these molecules form various types of ecDNA, including 1-LTR and 2-LTR full-length circles, as well as rearranged forms. These distinct structures suggest that ecDNA is not merely a passive byproduct, but the result of regulated molecular processes that intercept transposon replication intermediates. These circular molecules are abundantly detected from whole testes without any enrichment, suggesting that ecDNA formation is not accidental but a regulated process that actively diverts integration-competent DNA from reinserting into the genome. Given that retrotransposon integration can cause DNA damage and sterility, ecDNA formation likely serves as a buffering system, neutralizing transposon-derived DNA and preventing insertional mutagenesis. The absence of heritable new insertions in offspring further supports this conclusion.

Our study provides evidence that *Drosophila* spermatogenesis employs a novel protective mechanism: the sequestration of reverse-transcribed retrotransposon DNA into ecDNA, thereby preventing its reintegration into the genome. The formation of ecDNA as a genome defense mechanism raises new questions about how widespread and conserved this strategy may be. One intriguing direction for future research is to examine whether similar phenomena occur in other biological systems where retrotransposons are active but do not lead to significant genome integration. For instance, retrotransposon activation plays a critical role in mammalian embryogenesis(Macfarlan et al. 2012; Wang et al. 2014; Grow et al. 2015), yet unchecked integration during this period could introduce mutations that propagate through differentiating cell lineages and contribute to disease (Gerdes et al. 2016; DiRusso and Clark 2023). It is plausible that mammals, like *Drosophila*, have evolved similar strategies to harness retrotransposon activity for developmental processes while minimizing the risk of genomic insertion. Studying these contexts could reveal whether ecDNA formation is a common cellular strategy to manage transposon activation without compromising genome integrity.

## Materials and Methods

### Fly husbandry and strains

Flies were maintained at 25°C on standard *Drosophila* media containing molasses, enriched cornmeal, and inactivated yeast. The following fly stocks were used: sh-*white* (Bloomington Drosophila Stock Center #33762), sh-*aub* and sh-*ago3* (Senti et al., 2015), MTD*-*Gal4 (Bloomington Drosophila Stock Center #31777), *w^1118^*(Bloomington Drosophila Stock Center #5905), *rhi^2^* and *rhi^KG^* (Dr. William Theurkauf Lab).

### Fertility assay

A single one-day-old male virgin fly (MTD*-*Gal4>sh-*white* or MTD-Gal4>sh-*aub;ago3*) was crossed with 3 three-day-old female wild-type flies (*w^1118^*). After 24 hours, the male was crossed with 3 new three-day-old virgin flies and the original 3 females were flipped to fresh food every other day. This was repeated until the males were six days old. The number of eclosed flies from each cross was counted two weeks after the initial mating of each male with 3 virgin females. This assay was done with three replicates per genotype.

### RNA sequencing

Total RNA was extracted from fly testes and used as input for the TruSeq Stranded Total RNA Library Prep kit (Illumina). First, rRNA was depleted enzymatically with the Illumina Ribo-Zero Plus rRNA Depletion kit (Illumina Cat. # 20040526). Then, any residual DNA was removed by TURBO DNase (Invitrogen Cat. # AM2238) treatment for 30 minutes at 37°C and column purified with the RNA Clean and Concentrator-5 kit (Zymo Research Cat. # R1015) to remove the DNase enzyme. The purified RNA was then fragmented, the first strand of cDNA was synthesized, and the cDNA was purified with AMPure XP beads (Beckman Coulter Cat. # A63881). The RNA template was removed with RNAse H (Thermo Fisher Scientific Cat. # 18021014) and the second strand of cDNA was synthesized by DNA Polymerase I (NEB Cat. # M0210L) with dUTP incorporation. The DNA was purified with AMPure XP beads. Then the DNA ends were repaired, A tailing was added to the 3’ end of each strand, and adapters were ligated to the cDNA with bead purification performed after each of those steps. Then the cDNA was UDG treated and PCR amplified using Phusion High-Fidelity DNA Polymerase (NEB Cat. # M0530L). The libraries were sequenced through Azenta.

RNA-seq reads were mapped to *Drosophila* transposon consensus sequences with Bowtie2. Transposon reads were quantified by StringTie with the--rf option to indicate strandedness.

### ecDNA sequencing

For ecDNA sequencing, total DNA was extracted from testes using the Quick-gDNA MicroPrep Kit (Zymo Research Cat. # D3021). One µg of DNA was treated with Plasmid Safe DNase (Lucigen Cat.# E3110K) overnight at 37°C to remove linear DNA and then the enzyme was heat inactivated at 70°C for 30 minutes. Then the remaining circular DNA was purified with AMPure XP beads (Beckman Coulter Cat. # A63881). Then 5 ng of purified circular DNA was amplified though rolling circle amplification (RCA) using Phi29 polymerase (NEB Cat.# M0269L). RCA was carried out at 30°C for 18 hours and then the enzyme was heat inactivated at 65°C for 10 minutes. This yields a long piece of DNA with the contents of the original circular DNA repeated tandemly. Because this RCA product has several DNA branches that make it difficult to pass through the a nanopore for sequencing, the DNA is then debranched using T7 endonuclease I (NEB Cat. # 0302L). The short debranched fragments were then removed using the Short Read Eliminator XL buffer (Circulomics Cat. # SS-100-111-01) at room temperature overnight. The DNA was then purified through ethanol precipitation. The resulting DNA was used for library preparation using the standard Ligation Sequencing Kit V14 (ONT SQK-LSK114) protocol and sequenced using an R10.4.1 flow cell on GridION.

### Divergent PCR

For divergent PCR, total DNA was first extracted from the testes using the Zymo Quick-gDNA MicroPrep Kit (Zymo Research Cat. # D3021). Then 300 ng of DNA was used in a 100 µL reaction with Plasmid Safe DNase (Lucigen Cat. # E3110K). Linear DNA digestion was done overnight at 37°C to remove linear DNA and then the enzyme was heat inactivated at 70°C for 30 minutes. Then, 2 µL of digested DNA was directly used for PCR. CloneAmp HiFi PCR Premix (Takara Cat. # 639298) was used for PCR with the standard thermocycling conditions recommended by Takara. Since the mitochondrial DNA was present in lower abundance than the circular DNA, for diverent PCR from sh-*white* and sh-*aub;ago3* testes, the reaction had 40 cycles for the mitochondrial DNA and 35 cycles for the transposon DNA. For divergent PCR from *rhi* mutant tests, 35 cycles were used for both mitochondrial DNA detection and *nomad* circular DNA detection.

### RNA FISH

Stellaris RNA FISH probes for *nomad* and *copia* were synthesized by LGC Biosearch Technologies. Both the *nomad* and *copia* RNA FISH probe sets were labeled with Quasar 570 dye.

For RNA FISH, the testes were dissected in ice cold PBS and fixed in 4% formaldehyde diluted in PBS for 20 minutes at room temperature. The testes were washed once with PBST (PBS + 0.1% Triton X-100) and then twice with PBS, for five minutes each wash. Then the samples were permeabilized in 70% ethanol for 2 hours at 4°C. Then the testes were washed with Wash Buffer A (LGC Biosearch Technologies Cat. # SMF-WA1–60). The testes were hybridized with 0.5 µL of 12.5 µM transposon probe stock in 50 µL of hybridization buffer (LGC Biosearch Technologies Cat. # SMF-HB1–10) overnight at 37°C. Then the testes were washed twice more with Wash Buffer A for 30 minutes each at 37°C, with DAPI added at a 1:1000 dilution in the second wash. Then the testes had a 5-minute wash in Wash Buffer B (LGC Biosearch Technologies Cat. # SMF-WB1–20) at room temperature. The testes were mounted with Vectasheild Mounting Medium and sealed with nail polish. The testes hybridized with RNA FISH probes were imaged with a Stellaris 8 confocal microscope using LAS X software.

### Immunostaining

The following antibodies were used for immunofluorescence staining in the testis: aub (1:50 dilution, Brennecke Lab), ago3 (1:50 dilution, Brennecke Lab), *nomad* Gag (1:500 dilution, generated for this study by LifeTein), γH2Av (1:400 dilution, DSHB Cat. # UNC93-5.2.1), anti-mouse Alexa 488 secondary antibody (1:400 dilution, Molecular Probes Cat. # A21202), anti-rabbit 568 secondary antibody (1:400 dilution, Invitrogen Cat. # A-11011).

The *nomad* Gag rabbit polyclonal antibody was generated by LifeTein. The lyophilized antibody was reconstituted in water at a concentration of 1 mg/mL. That reconstituted antibody was then used at a dilution of 1:500. As a control the lyophilized pre immune sera was also dissolved in water to a concentration of 1 mg/mL and used at a dilution of 1:500.

For immunofluorescence staining, the testes were dissected from flies in ice cold PBS and fixed in 4% PFA diluted in PBST for 20 minutes at room temperature. The tetes were then washed three times in PBST. Blocking was carried out with 10% normal goat serum (NGS) (Sigma-Aldrich Cat. # G9023) diluted in PBST for 1 hour at room temperature. The testes were incubated with primary antibody in blocking solution (10% NGS) overnight at 4°C. The samples were then washed three times in PBST and incubated with secondary antibody diluted in blocking solution for 2 hours at room temperature.

Then samples were washed three more times in PBST, with DAPI (1:1000, Abcam Cat. # ab228549) in the second wash step. Finally, the samples were mounted with Vectashield Mounting Medium. Images were taken on a Leica Stellaris 8 confocal microscope with LAS X software.

### Genomic DNA sequencing (carcass and testes)

Testes and carcass were dissected in cold PBS. The carcass includes the entire fly body without the male reproductive tract. Genomic DNA was extracted from the samples using Zymo Quick DNA extraction kit (Zymo Research Cat. # D3021) and the quality of the DNA was assessed by Nanodrop. For the carcass DNA, the sequencing library was prepared according to the Nanopore Rapid Sequencing gDNA-barcoding (ONT SQK-RBK004) protocol. The barcoded samples were pooled and sequenced on a single flow cell. The testes genomic sequencing libraries were prepared following the standard Nanopore Rapid sequencing DNA – PCR barcoding (SQK-RPB114) protocol. The barcoded testes gDNA samples were also pooled and sequenced on a single separate flow cell.

### Genomic DNA sequencing (larvae)

Single crosses of one male (MTD-Gal4>sh-*white* or MTD-Gal4>sh-*aub;ago3*) were crossed with a single virgin wild type female (*w^1118^*). After larvae hatched, three larvae per cross were collected for DNA extraction. DNA was extracted from each parent and the offspring with the Zymo Quick DNA extraction kit (Zymo Research Cat. # D3021). The quality of the extracted DNA was assessed by Nanodrop and used as input for the Nanopore Rapid Sequencing gDNA-barcoding (ONT SQK-RBK004) kit following the standard protocol. The samples were pooled and sequenced on a single flow cell using GridION.

### ecDNA sequencing analysis

For the ecDNA-seq data, the adapter sequences were trimmed from the reads and any read with adapter sequences in the middle of the read were discarded. The trimming was done with porechop. Then the trimmed reads were mapped to the *Drosophila* transposon consensus sequences using minimap2 with the following parameters:-ax map-ont-Y-t 16. Reads were classified as circular DNA if the read had multiple alignments to the transposon and those alignments were directly adjacent, withing 100 bp of each other, or overlapping. The reads were then classified into different circle types. The 1-LTR full-length circles were named when the read fully mapped to the internal sequence of the transposon, but the portion of the reads mapping to the end sequences overlapped by the length of the LTR. The reads were designated as 2-LTR full-length if the read fully mapped to the transposon sequences and none of the subsequent alignments overlapped with each other. The reads were designated as 1-LTR rearranged if the read aligns to one end of the transposon fully covering the LTR, but does not map to the other LTR. The reads are classified as non-LTR rearranged when the read maps to the internal transposon sequence and does not fully cover either LTR. The coverage for each circle type was plotted in R using the circos function of the circlize package. The coverage was determined by using the bedtools genomecov function.

### Transposon copy analysis from genomic DNA

Fastq files were down sampled using rasusa to ensure equal coverage between samples that were being compared. The reads from the down sampled fastq files were then trimmed with porechop (and reads with middle adapters were removed) and then aligned to the *Drosophila* dm6 genome using minimap2 with the parameters:-ax map-ont-Y-t 16. For standard detection of non-reference transposon copies, the TLDR pipeline was used. For TLDR analysis, the standard requirement of 3 reads spanning each transposon copy were needed to call as a non-refence copy.

For one end reads, the genomic data was mapped to the consensus transposon sequences and *Drosophila* dm6. Reads containing 1 kb + of either end of the transposon plus 200 kb of flanking sequence were analyzed. The flanking DNA was then intersected with repeat sequences from a bed file of the repeat masker masked sequences for *Drosophila* dm6 using bedtools intersect-wao. If the flanking sequence was from the same transposon it was kept as a potential circular DNA read. If there was at least 200 kb of unique mapping sequence, the read was retained as unique flanking sequence. Reads with unique flanking sequence were retained and further analyzed to determine how many potential *nomad* copies could be detected by one end reads. One end reads with location sites within 2 kb of the genome were retained as the same copy. One end transposon reads entirely flanked by repetitive sequences were discarded as they could or accurately be assigned a genomic location.

## Acknowledgements

We thank Drs. William Theurkauf and Julius Brennecke and Bloomington *Drosophila* Stock Center for providing fly alleles. We thank members from ZZ lab for constructive suggestions during the project development. This work was supported by the grants to Z.Z. from the Pew Biomedical Scholars Program, the Sontag Distinguished Scientist Award program, and the National Institutes of Health (R01 GM141018 and R01 GM152423).

## Author Contributions

Z.Z. and L.T. conceived the project and performed data analysis. L.T. performed all experiments. Z.Z. and L.T. wrote the manuscript.

---

**Supplemental Figure 1.**
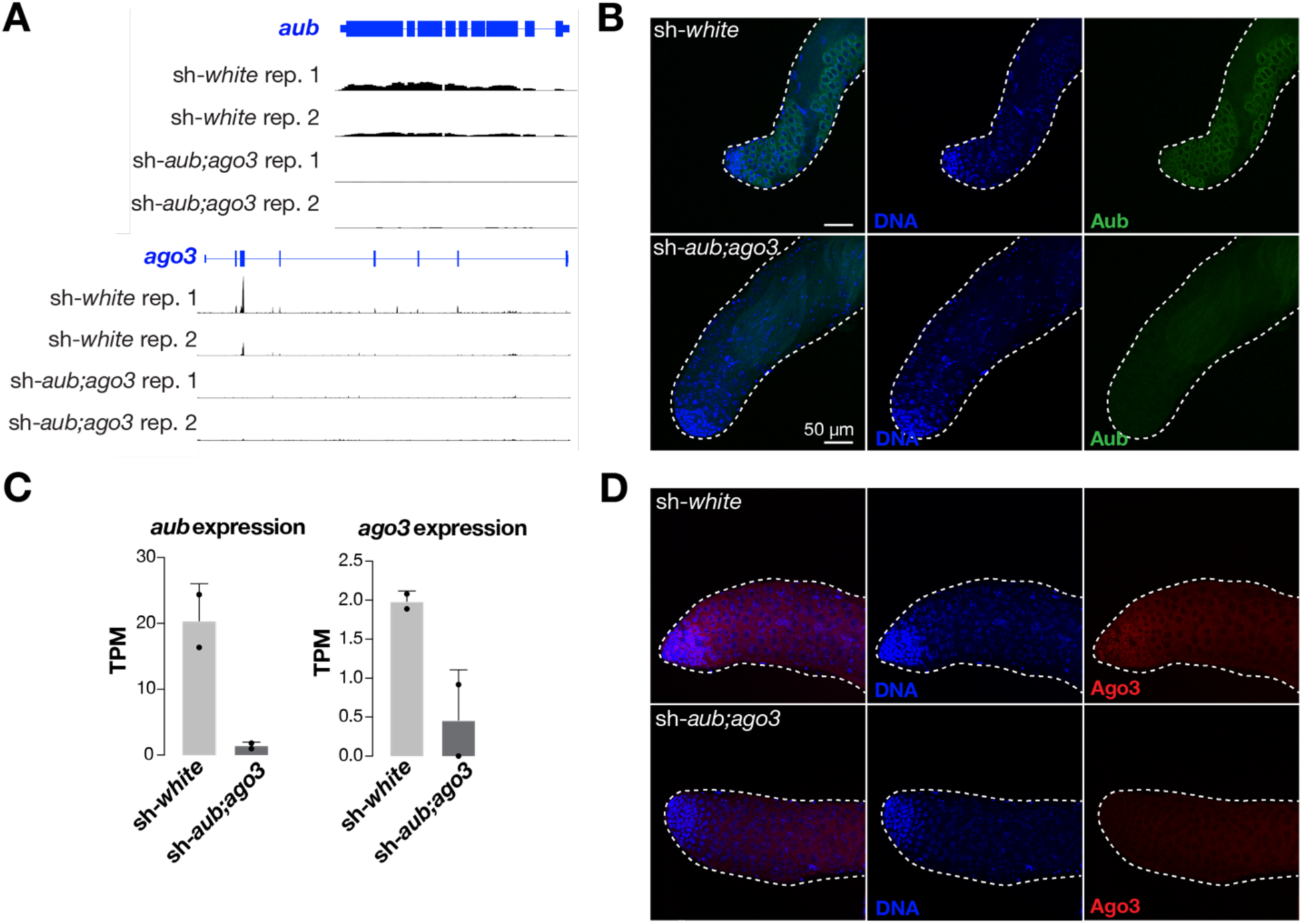
Aub and Ago3 are efficiently depleted by RNAi in the testis. **A.** UCSC browser display of *aubergine* and *argounaute3* RNA expression from replicates of sh-*white* and sh-*aub;ago3* testes. **B.** Quantification of *aubergine* and *argonaute3* RNA expression from RNA sequencing. **C.** Immunofluorescence staining for Aub and Ago3 at the apical tip of testes dissected from 3-day-old flies. The sh-*aub* and sh-*ago3* constructs in these flies are driven by MTD-Gal4.

**Supplemental Figure 2.**
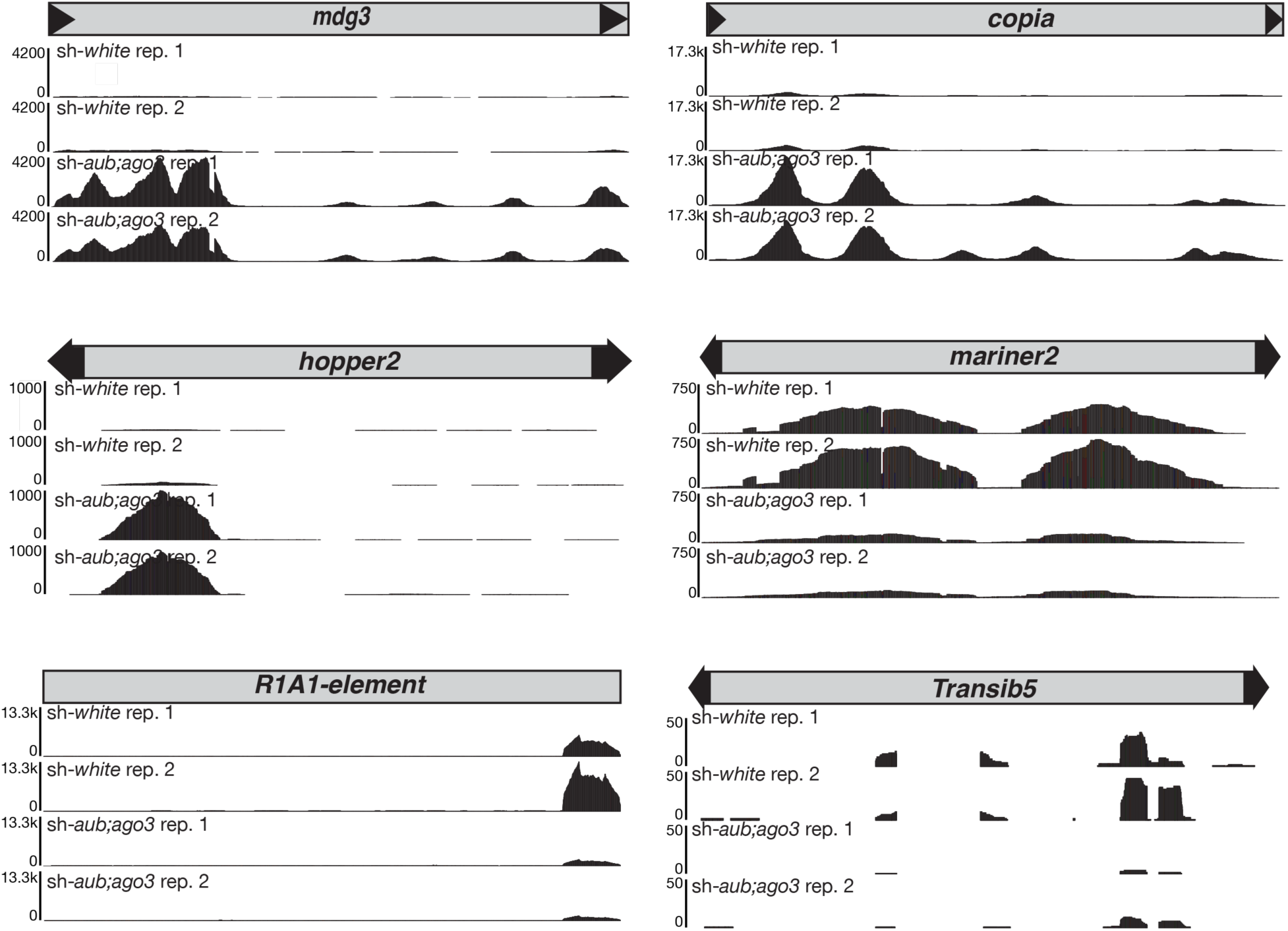
Visualization of RNA-seq reads for transposons that are differentially expressed in control and transposon derepressed testes. RNA-seq reads from across the length of each transposon are visualized using IGV.

**Supplemental Figure 3.**
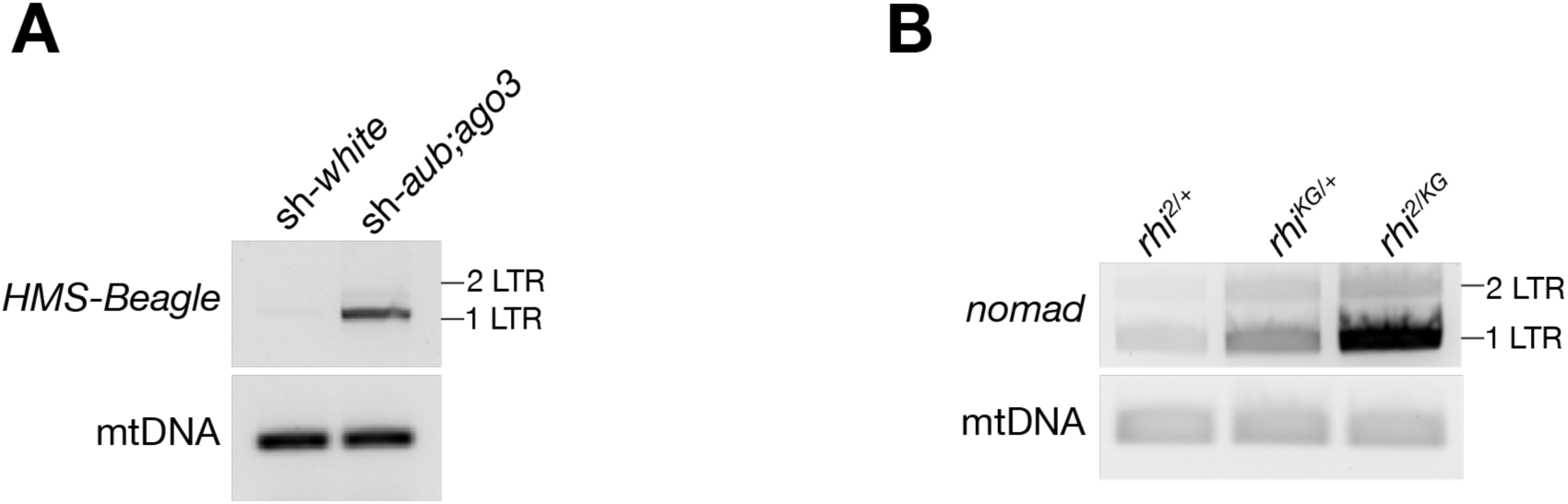
Transposon ecDNA can be detected by divergent PCR. A. Divergent PCR for *HMS-Beagle* from control and transposon derepressed testes. Input for PCR was DNA from 3-day-old testes with linear DNA digested. **B.** Divergent PCR for *nomad* ecDNA from 1-day-old testes of heterozygous control testes or *rhino* mutant testes. Linear DNA was digested from the sample.

**Supplemental Figure 4.**
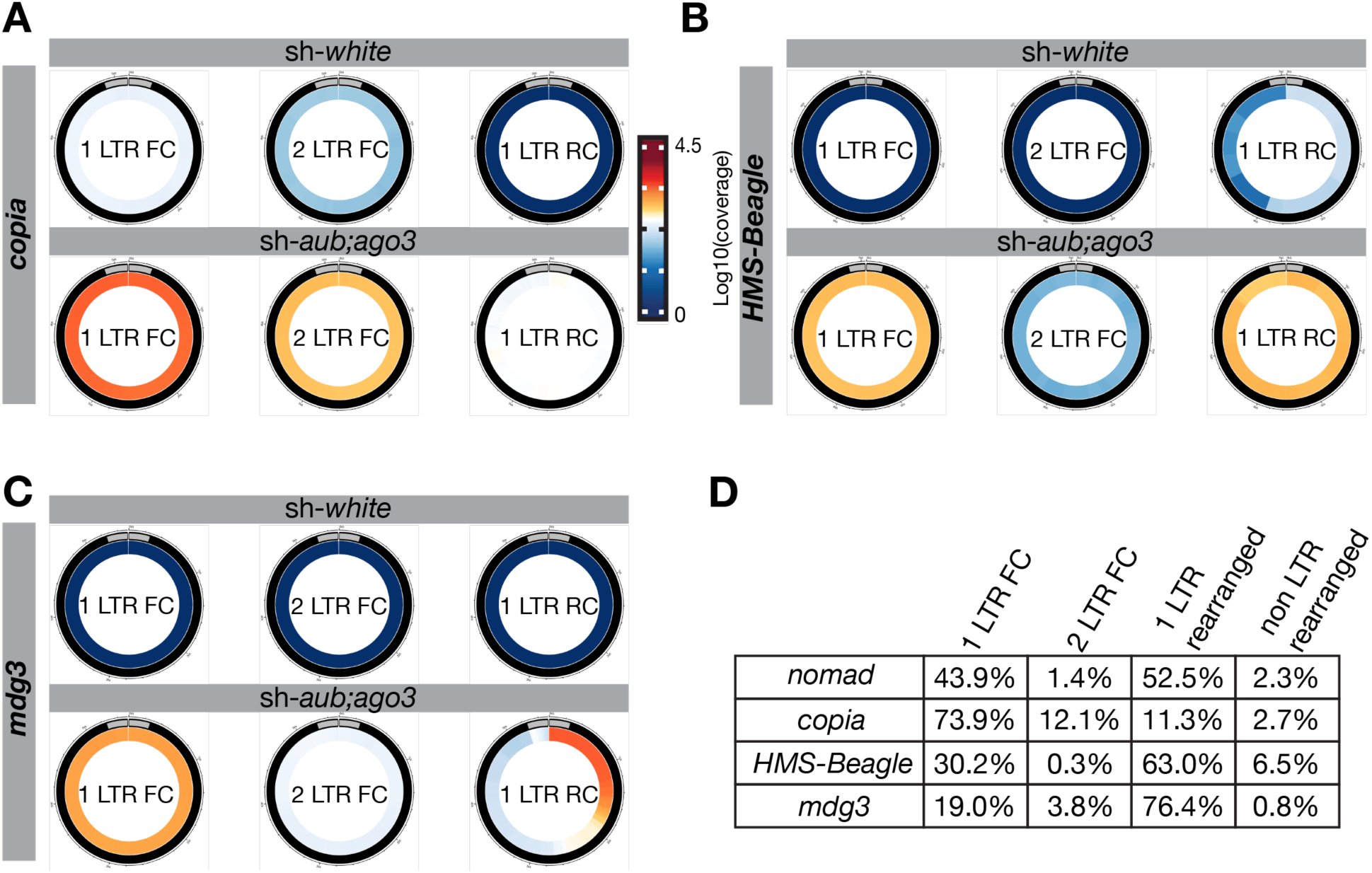
LTR retrotransposons predominantly make 1-LTR ecDNA. A-C. Circos plots of different types of LTR retrotransposon ecDNA and their abundance in testes from 3-day-old flies. Plots are from one replicate of ecDNA sequencing, each with a similar amount of mitochondrial DNA. Gray boxes signify the LTR region of the retrotransposon. FC = full-length circle. RC = rearranged circle. **D.** Chart reporting the percentage of ecDNA circles that correspond to each type of circle from sh-*aub;ago3* testes ecDNA sequencing. This is average data from three replicates of sh-*aub;ago3* ecDNA sequencing. Each replicate was normalized by the mitochondrial DNA content from that sample.

**Supplemental Figure 5.**
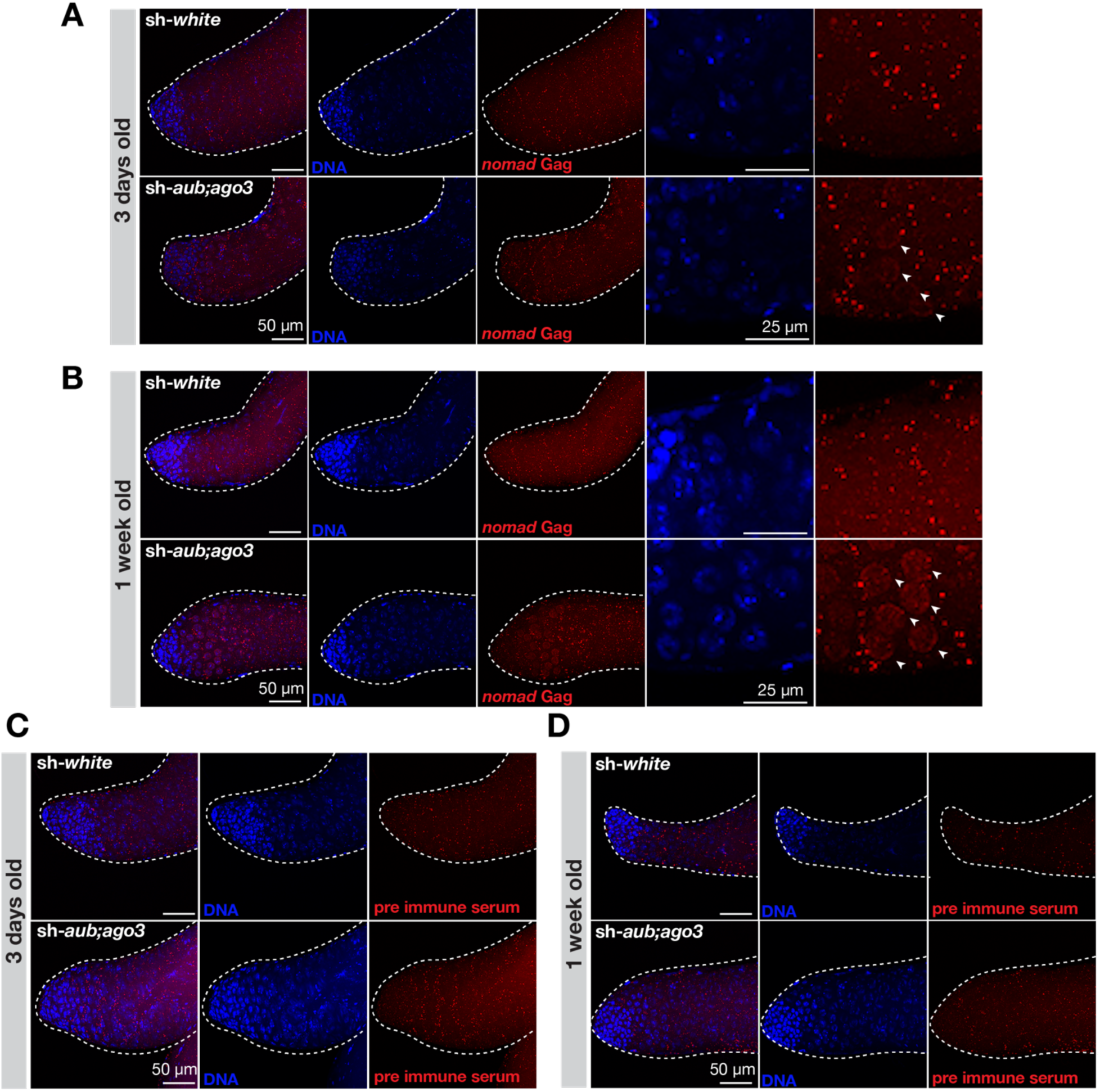
n*o*mad Gag is produced in spermatocytes. A. *nomad* Gag immunostaining at the apical tip of testes dissected from 3-day-old flies. **B.** *nomad* Gag immunostaining at the apical tip of testes dissected from 1-week-old flies. **C.** Three-day-old testes incubated with pre immune serum in replacement of *nomad* Gag antibody. **D.** Testes from 1-week-old flies incubated with pre immune serum in replacement of *nomad* Gag antibody.

**Supplemental Figure 6.**
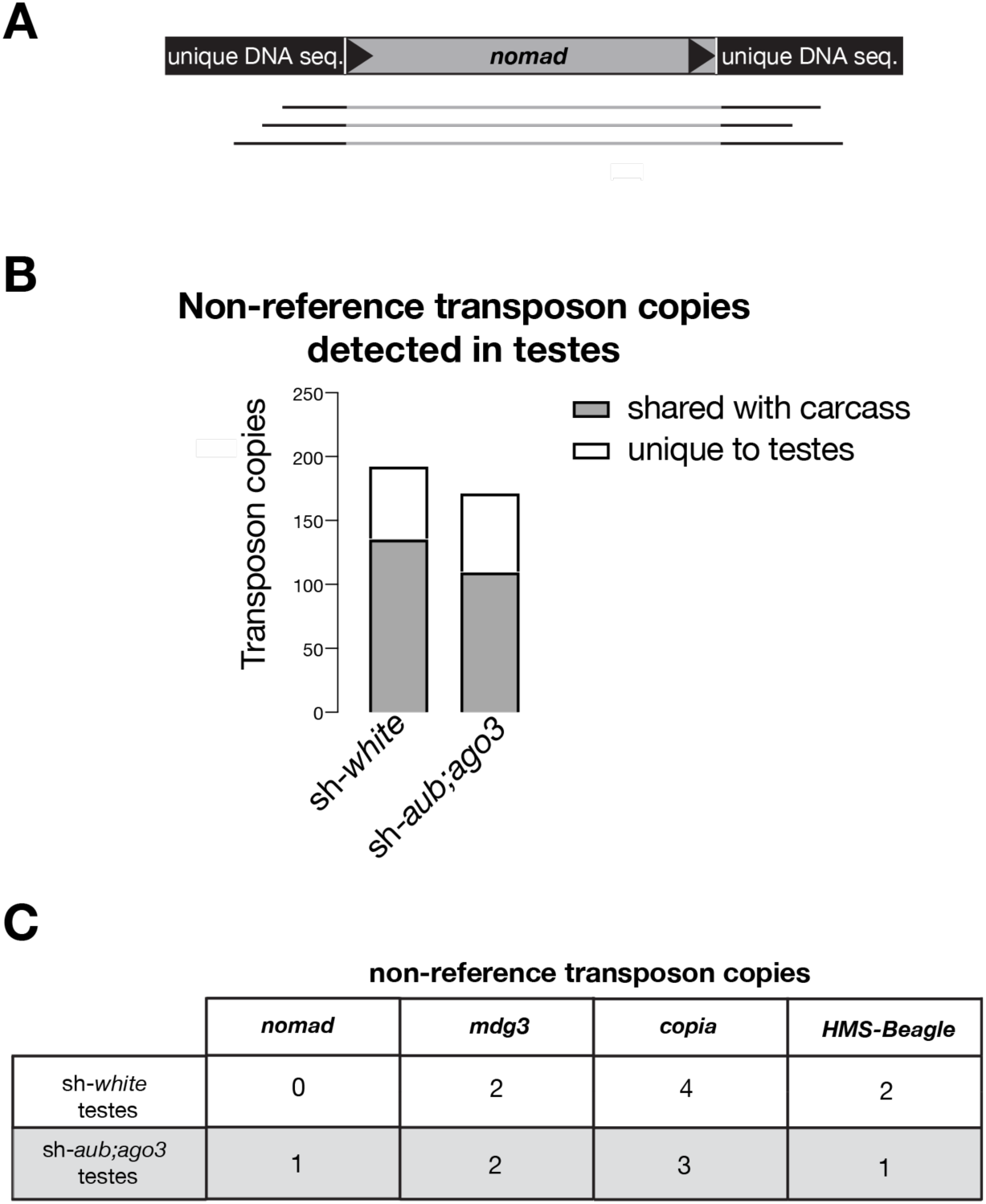
L**T**R **retrotransposons that make ecDNA in the testes rarely integrate. A.** Model of the types of reads that were used for panels B and C. **B.** Total transposon copies detected by the TLDR pipeline in the testes of control and transposon derepressed flies. This pipeline requires three or more reads to support each copy with flanking sequence. **C.** Transposon copies detected in sh-*white* and sh-*aub;ago3* testes for LTR retrotransposons that make significantly more ecDNA upon derepressing transposons.

**Supplemental Figure 7.**
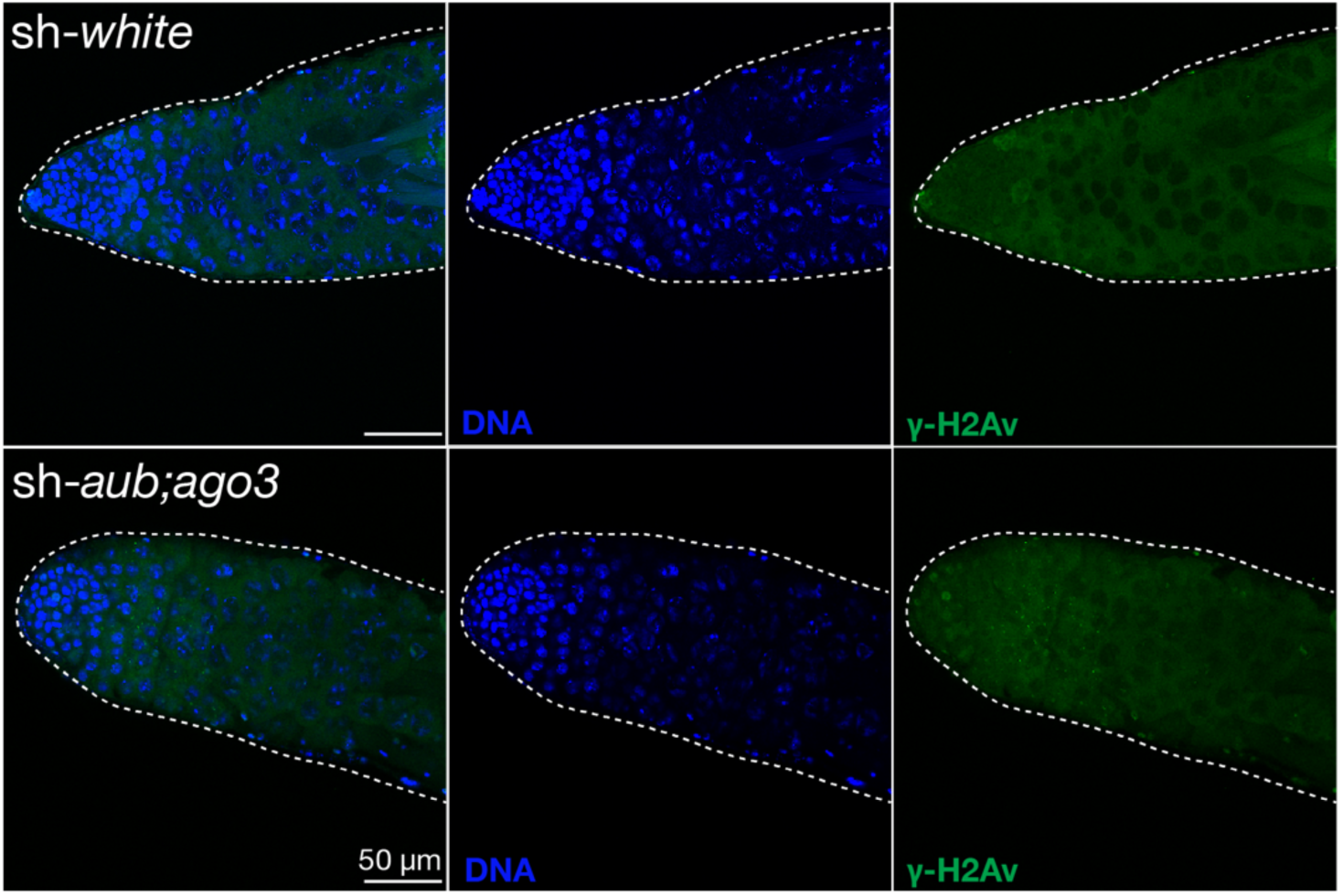
There is minimal DNA damage in transposon derepressed testes. γ-H2Av immunostaining for DNA double-strand breaks at the apical tip of testes dissected from 3-day-old flies.

**Supplemental Figure 8.**
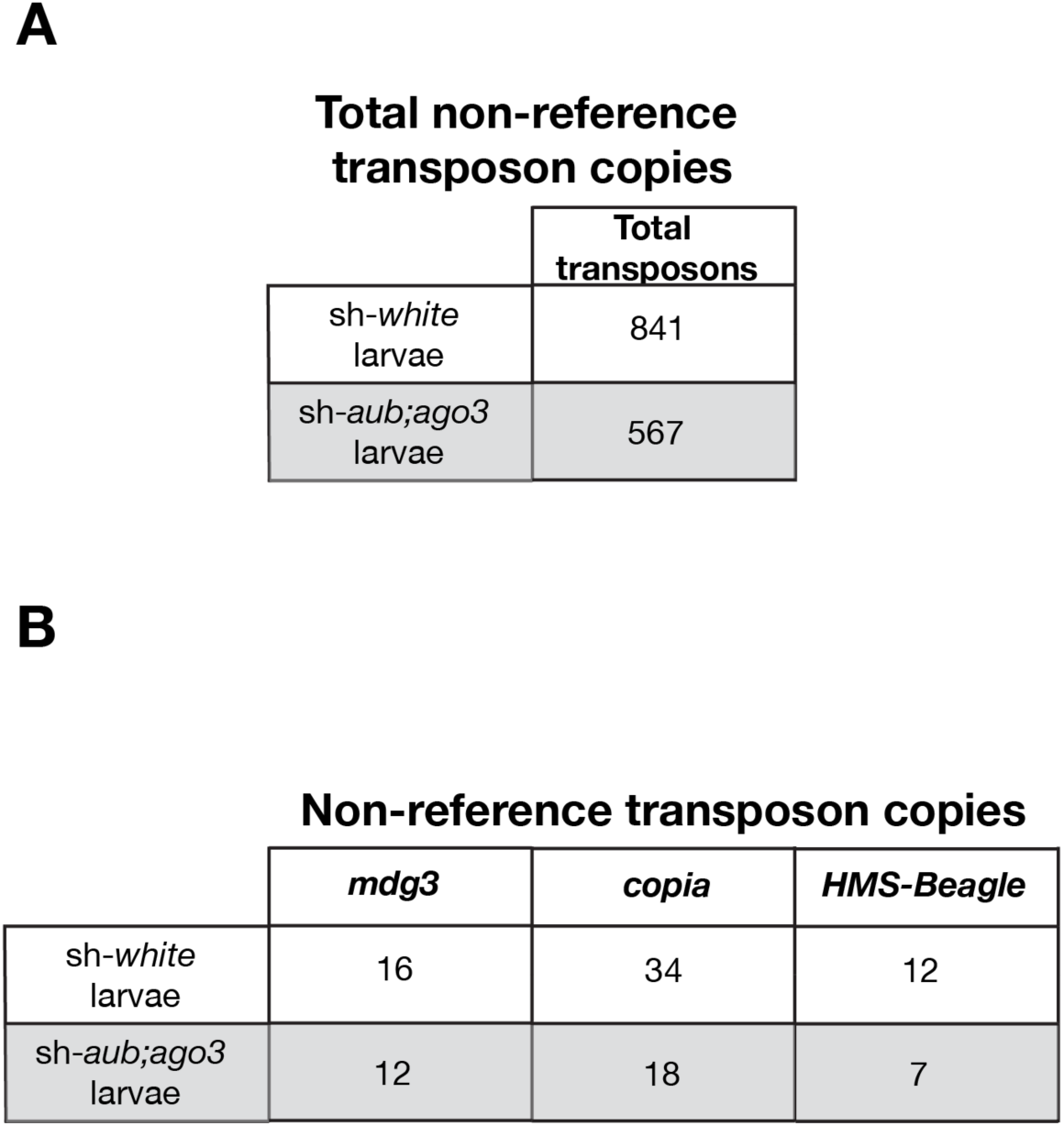
O**f**fspring **from males with transposons derepressed in the germline do not have an increase in transposon genomic transposon copies. A.** Total non-reference transposon copies detected in the offspring from crosses of wild-type females crossed with either a control male (MTD-Gal4>sh-*white*) or male with transposons derepressed (MTD-Gal4>sh-*aub;ago3*) in the germline. Transposons have at least 3 reads supporting each copy and flanking sequence. **B.** Non-reference transposon copies from LTR retrotransposons that make ecDNA.

## Supplemental Material Description

**Supplemental Figure S1**-This figure shows that two primary components of the piRNA pathway, Aub and Ago3, are able to be depleted by RNAi. This is foundational for the paper because this is the model of transposon de-repression used throughout the paper.

**Supplemental Figure S2**-This figure shows a more detailed view of how disrupting the piRNA pathway can change transposon RNA expression.

**Supplemental Figure S3**-This figure shows that transposon ecDNA can be detected by PCR in two different genetic models of transposon activation (depleting Aub and Ago3 by RNAi/ mutating *rhino*).

**Supplemental Figure S4**-This figure supplements a main conclusion that LTR retrotransposon ecDNA can integrate into itself (shown by 1-LTR rearranged circle coverage plots). It also shows the proportion of each circle type that is made by other transposons in addition to *nomad*.

**Supplemental Figure S5**-This figure supports the conclusion that *nomad* products ae produced in the spermatocytes by showing that *nomad* Gag protein is produced in those cells upon Aub/Ago3 depletion.

**Supplemental Figure S6**-This figure supports the conclusion that transposons do not abundantly integrate into the genomes of testes cells upon Aub/Ago3 depletion.

**Supplemental Figure S7**-This figure supports the conclusion that transposition is rare in the testes by showing minimal DNA double-strand breaks in the testes of both sh-*white* and sh-*aub;ago3* testes.

**Supplemental Figure S8**-This figure supports the conclusion that transposition is rare in the testes even when transposons are derepresses, as there is not an increase in transposon content in the offspring of sh-*aub;ago3* males.

---

**Supplemental Table S1.**
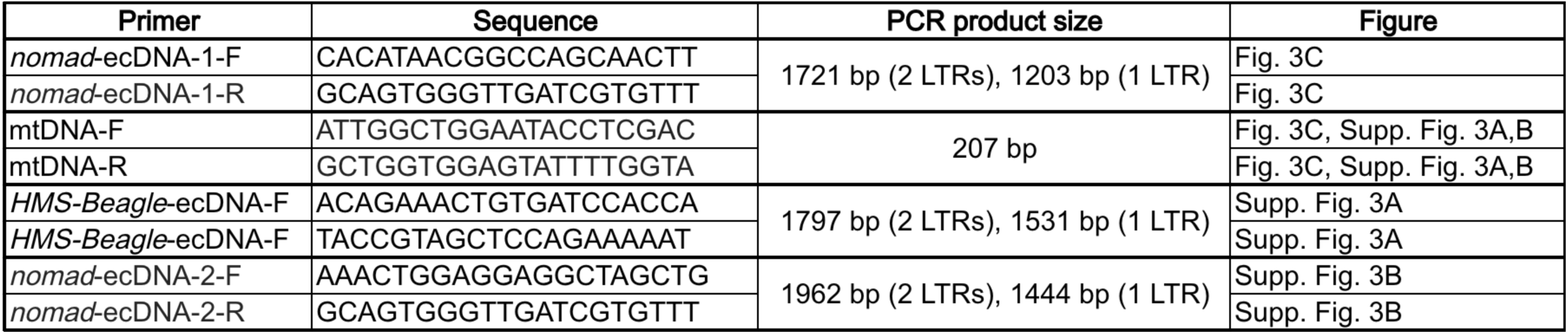
This table reports the primers used for divergent PCR for Fig. 3 and Supplemental Fig. 3.

**Supplemental Table S2.**
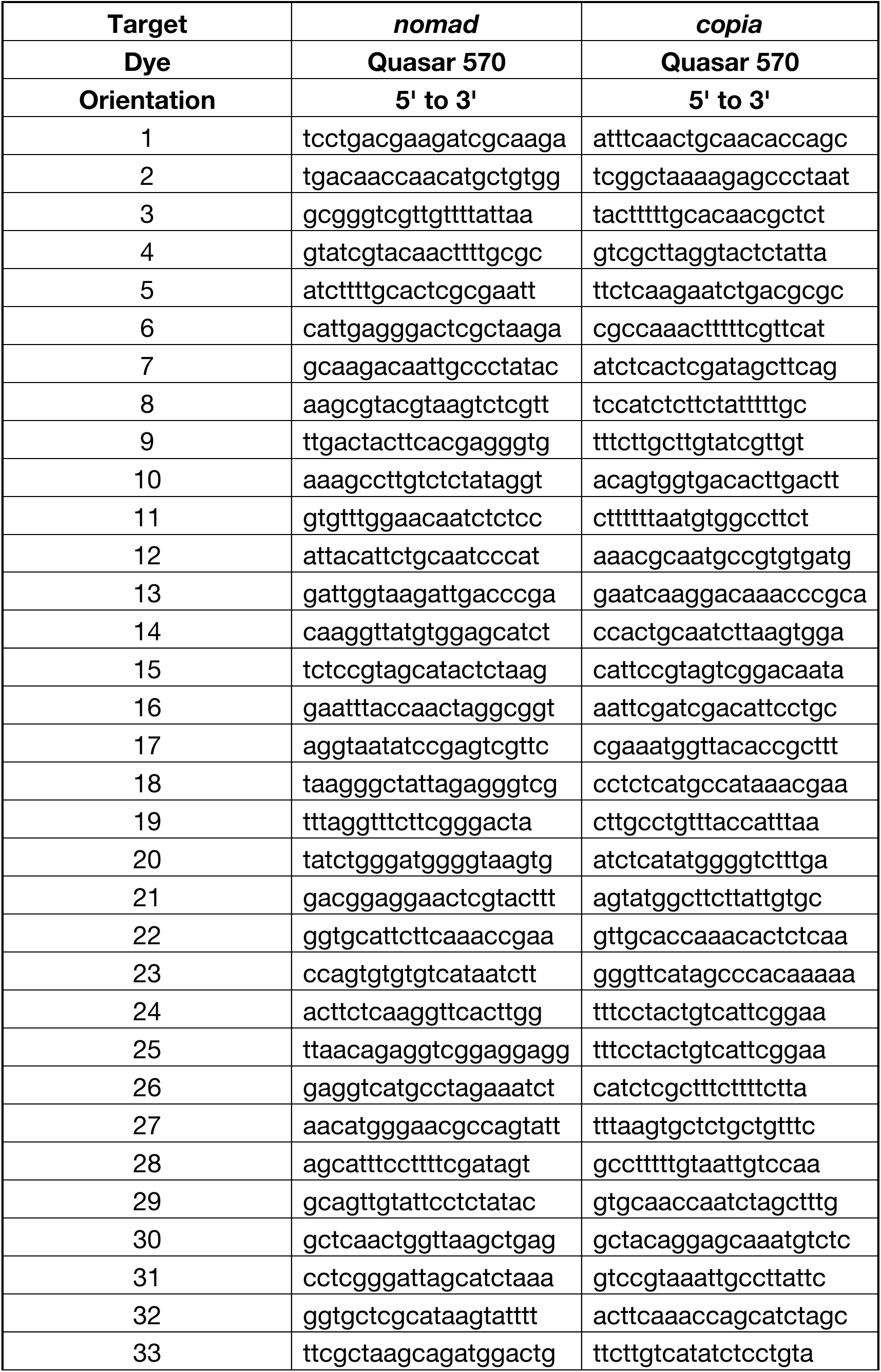
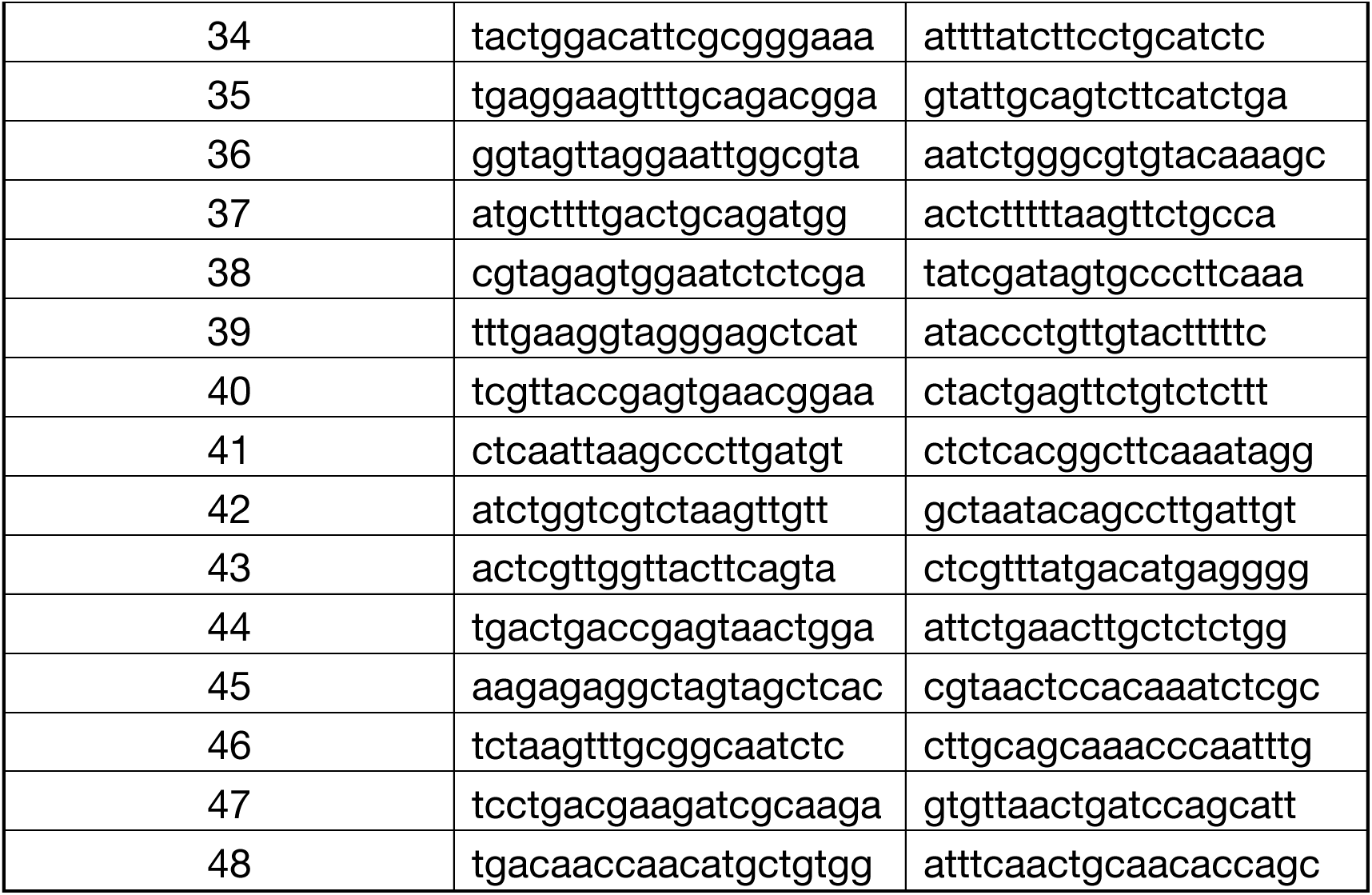
This table reports the RNA FISH probe sets for *nomad* and *copia* used in Fig. 4.

**Supplemental Table S3.** This table provides basic information (library preparation kit, read statistics, etc.) about the sequencing data used in this manuscript.

RNA Sequencing
| Sample | Barcode sequence | Number of reads | Total bases (Mbases) | Mean quality score | %Bases >= 30 |
| --- | --- | --- | --- | --- | --- |
| mtdGal4>sh-*white* | TTAGGC | 38,837,960 | 11,651 | 35.17 | 90.03 |
| mtdGal4>sh-*white* | TGACCA | 114,176,869 | 34,253 | 35.6 | 91.96 |
| mtdGal4>sh-*aub;ago3* | ACTTGA | 54,148,304 | 16,244 | 35.63 | 92.16 |
| mtdGal4>sh-*aub;ago3* | CAGATC | 63,803,380 | 19,141 | 35.68 | 92.39 |

ecDNA Sequencing
| Sample | Barcode | Sequencing Kit | Number of reads | Read length N50 | Median read length | Mean read length | Total bases |
| --- | --- | --- | --- | --- | --- | --- | --- |
| sh-*white* ecDNA rep. 1 | barcode16 | SQK-LSK114 | 106128 | 13475 | 2168 | 5505 | 584269829 |
| sh-*white* ecDNA rep. 2 | barcode17 | SQK-LSK114 | 93700 | 15458 | 4915 | 8032 | 752688230 |
| sh-*white* ecDNA rep. 3 | barcode18 | SQK-LSK114 | 101502 | 15021 | 5417 | 8173 | 829610849 |
| sh-*aub;ago3* ecDNA rep. 1 | barcode19 | SQK-LSK114 | 157943 | 11751 | 1545 | 4432 | 700106416 |
| sh-*aub;ago3* ecDNA rep. 2 | barcode20 | SQK-LSK114 | 180730 | 13496 | 1492 | 4777 | 863412071 |
| sh-*aub;ago3* ecDNA rep. 3 | barcode21 | SQK-LSK114 | 220033 | 12570 | 2036 | 5333 | 1173626493 |

Genomic DNA Sequencing
| Sample | Barcode | Sequencing Kit | Number of reads | Read length N50 | Median read length | Mean read length | Total bases |
| --- | --- | --- | --- | --- | --- | --- | --- |
| sh-*white* carcass | barcode02 | SQK-RBK004 | 416487 | 6700 | 2786 | 4034 | 1680488921 |
| sh-*aub;ago3* carcass | barcode04 | SQK-RBK004 | 243014 | 12295 | 4555 | 7005 | 1702390396 |
| sh-*white* testes | barcode05 | SQK-RPB114 | 1914773 | 5393 | 4776 | 4879 | 9343693083 |
| sh-*aub;ago3* testes | barcode06 | SQK-RPB114 | 1842002 | 4772 | 4038 | 4227 | 7786804192 |
| Offspring of sh-*white* males | barcode09 | SQK-RBK004 | 454643 | 10529 | 3682 | 5869 | 2668694551 |
| Offspring of sh-*aub;ago3* males | barcode12 | SQK-RBK004 | 467828 | 7382 | 2812 | 4350 | 2035184544 |

